# Convergent pathways with impaired inhibition at the frontal cortex are the outcome of differential alterations in the male and female schizophrenia model

**DOI:** 10.64898/2025.12.04.692295

**Authors:** Blanca Sánchez-Moreno, Alberto Mesa-Lombardo, María de la Fuente-Fernández, Ana Isabel Fraga-Sánchez, Ángela Calzado-González, Inés García-Ortiz, Miriam Martínez-Jiménez, Jorge García-Piqueras, David Vega-Avelaira, Claudio Toma, Ángel Núñez, Javier Gilabert-Juan

**Affiliations:** Department of Anatomy, Histology and Neuroscience, School of Medicine, Universidad Autonoma de Madrid, Madrid, Spain. Ph.D. Program in Neuroscience Cajal-UAM, School of Medicine, Universidad Autonoma de Madrid, Madrid, Spain; Centre for Molecular Biology Severo Ochoa (CBMSO), The Spanish National Research Council (CSIC) & Autonomous University of Madrid, Madrid, Spain; Neuroscience Research Australia, Sydney, NSW, Australia; School of Clinical Medicine, Discipline of Psychiatry and Mental Health, University of New South Wales, Sydney, NSW, Australia

**Keywords:** double-hit schizophrenia model, GABAA, GABAB, plasticity, inhibitory neurotransmission, parvalbumin

## Abstract

Schizophrenia is associated with impaired inhibitory neurotransmission and disrupted synaptic plasticity in the medial prefrontal cortex (mPFC), yet the biological mechanisms underlying these deficits may differ between sexes. Here, we used a double-hit rat model, combining perinatal NMDA receptor blockade and post-weaning social isolation, to dissect sex-specific alterations in inhibitory circuit maturation, synaptic plasticity, and prefrontal function. Male double-hit rats exhibited robust schizophrenia-like behaviors, reduced parvalbumin (PV), OTX2, and perineuronal net (PNN) expression, decreased GAD67 levels, and increased DNA damage in PV interneurons, together indicating impaired inhibitory maturation and weakened plasticity. In contrast, females showed milder behavioral deficits but displayed increased PV and OTX2 intensities, enhanced cFos activation in excitatory neurons, and transcriptomic upregulation of glutamatergic, GABAergic, and synapse assembly pathways, suggesting a state of heightened or dysregulated plasticity. Despite these divergent molecular trajectories, in vivo electrophysiology revealed a shared functional endpoint in both sexes: a shift from paired-pulse inhibition to facilitation during basolateral amygdala–evoked responses, reflecting impaired GABAB-mediated inhibitory feedback. Pharmacological blockade of GABAB, but not GABAA, receptors reproduced this phenotype, identifying GABAB signaling as a key mechanism underlying cortical disinhibition. Altogether, our findings reveal that males and females reach convergent prefrontal inhibitory deficits through sex-specific molecular pathways, underscoring the importance of sex as a biological variable in the pathophysiology and treatment of schizophrenia.

## Introduction

Schizophrenia is a heterogeneous neuropsychiatric disorder marked by disturbances in perception, motivation, and cognition (Tandon et al., 2024). Decades of research have not fully elucidated its pathophysiology. A unifying framework is offered by the neurodevelopmental theory, which postulates that early-life disturbances in brain maturation, caused by genetic predisposition, perinatal stress, or other environmental insults, predispose people to the later onset of psychotic symptoms (Fatemi & Folsom, 2009; Schmitt et al., 2023). These developmental abnormalities are thought to gradually but continually change synaptogenesis, neuronal migration, and the building of brain circuits. When cortical and subcortical circuits undergo critical refinement throughout adolescence and early adulthood, these changes lead to vulnerabilities that persist into adulthood and cause the disease (Hall & Bray, 2022).

The evidence that these neurodevelopmental traumas cause long-lasting changes in synaptic plasticity is growing. Connectivity impairments and decreased dendritic spine density, especially in the prefrontal cortex, have all been documented in studies conducted on both human postmortem tissue and animal models (Banks et al., 2015; Hasan et al., 2011; Konopaske et al., 2014; Uhlhaas & Singer, 2010). These deficiencies in plasticity jeopardize experience-dependent circuit remodeling and are thought to underlie patients’ working memory and cognitive impairments (Dienel et al., 2022). OTX2 is a homeoprotein transcription factor critical for the maturation of parvalbumin-positive interneurons and the formation of perineuronal nets (PNNs) in the brain regions controlling these processes (Parmar et al., 2024; Sugiyama et al., 2008; Vincent et al., 2021). Disruptions in OTX2 signaling can delay or destabilize PNN assembly, thereby impairing the closure of critical periods of plasticity and contributing to persistent Excitatory/Inhibitory (E/I) imbalance (Lee et al., 2017). Such alterations may synergize with deficits in neurotrophic pathways, including mTOR and BDNF-TrkB signaling, further compromising interneuron function and preventing the proper stabilization and refinement of cortical circuits (Gibel-Russo et al., 2022; Parmar et al., 2024). In turn, OTX2 controls parvalbumin (PV) interneuron maturation, and dysfunction of PV interneurons and altered expression of GABAergic receptors disrupt the E/I balance, impairing network synchrony, gamma oscillations, and information processing, traits observed in patients of schizophrenia (Arazi et al., 2025; Beurdeley et al., 2012; Guyon et al., 2021). These disturbances are thought to interact with plasticity deficits, amplifying cortical dysconnectivity and cognitive dysfunction. Collectively, converging evidence supports a model in which early neurodevelopmental insults perturb both synaptic plasticity and E/I balance, producing a cascade of circuit-level dysfunction that contributes to the clinical phenotype of schizophrenia (Le Roux et al., 2008; Mana et al., 2024).

In addition, sex-related differences in schizophrenia extend beyond clinical presentation and suggest distinct neurobiological trajectories shaped by hormonal and molecular factors (Gogos et al., 2015; Mendrek & Mancini-Marïe, 2016). Epidemiological studies indicate that males typically exhibit earlier onset, greater prevalence of negative and cognitive symptoms, and more severe functional impairment, whereas females often show later onset and relatively preserved social and cognitive functioning (Fountoulakis et al., 2022; Rotstein et al., 2018; Zhang et al., 2025). At the mechanistic level, estrogens have been implicated in modulating synaptic plasticity, dendritic spine density, and neurotrophic signaling, potentially buffering against excitatory deficits and interneuron dysfunction (Gogos & van den Buuse, 2015; Woolley, 1998). Conversely, males appear more susceptible to disruptions in parvalbumin-positive interneuron maturation and GABAergic transmission, exacerbating E/I imbalance (Santos-Silva et al., 2024; Wang et al., 2025; Woodward & Coutellier, 2021). Postmortem and neuroimaging evidence further supports sex-specific alterations in the prefrontal region, consistent with divergent patterns of cortical dysconnectivity (Carceller et al., 2024; Iraji et al., 2022, 2024). Together, these findings suggest that plasticity and E/I regulation are differentially modulated by sex hormones and developmental timing, contributing to the heterogeneity of symptom expression and disease course across sexes. However, the molecular mechanisms that operate differentially in each sex are unknown, and few studies compare them.

In this context, it is imperative to dissect sex-specific modulators of schizophrenia pathogenesis and to specify which structural and functional features of inhibitory neurotransmission and synaptic plasticity diverge between males and females. To probe these mechanisms, the double-hit murine model, combining perinatal MK801 administration (an antagonist of NMDA receptors) and post-weaning social isolation, offers a robust paradigm to study deficits in inhibitory circuit development, plasticity, and E/I balance (Castillo-Gómez et al., 2017; Garcia-Mompo et al., 2020; Gilabert-Juan et al., 2013; Klimczak et al., 2024). In the present study, we will systematically compare male and female double-hit rats, aiming to delineate sex-dependent structural, molecular, and functional differences in the medial prefrontal cortex (mPFC), with a focus on interneuron plasticity.

## Materials and methods

### 1. Animals

Eleven pregnant Lister Hooded rats were purchased from Inotiv/Envigo and housed individually in a controlled temperature room and on a 12-h light/dark cycle with food and water available *ad libitum*. 49 female and 56 male pups were born from the pregnant rats and used for the experiments. Pups within each litter were randomly assigned to the double hit model group or to the control group.

At P7 rat pups were intraperitoneally injected either with a solution of the non-competitive antagonist of the N-methyl-D-aspartate (NMDA) receptor MK-801 (1 mg/kg, Merck) or saline solution (0.9% NaCl). All rat pups remained with their mothers until weaning. At P21 double-hit group rats were isolated in 220 x 220 x 145 mm cages, while control rats were socially housed in groups of three in 215 x 465 x 145 mm cages. All rats were housed in the same room, sharing the same controlled light, temperature, and humidity. Rats reared in isolation could hear and smell other rats but were unable to see or have physical contact with them. Rats were reared in these conditions for at least 10 weeks, until they reached adulthood (Figure 1A). MK-801 injection caused a significant reduction in weight in both female and male pups when compared to pups injected with saline, but this difference disappeared at P40 in males (Supplementary Figure S1A) and decreased at P40 in females (Supplementary Figure S1B).

**Figure 1:**
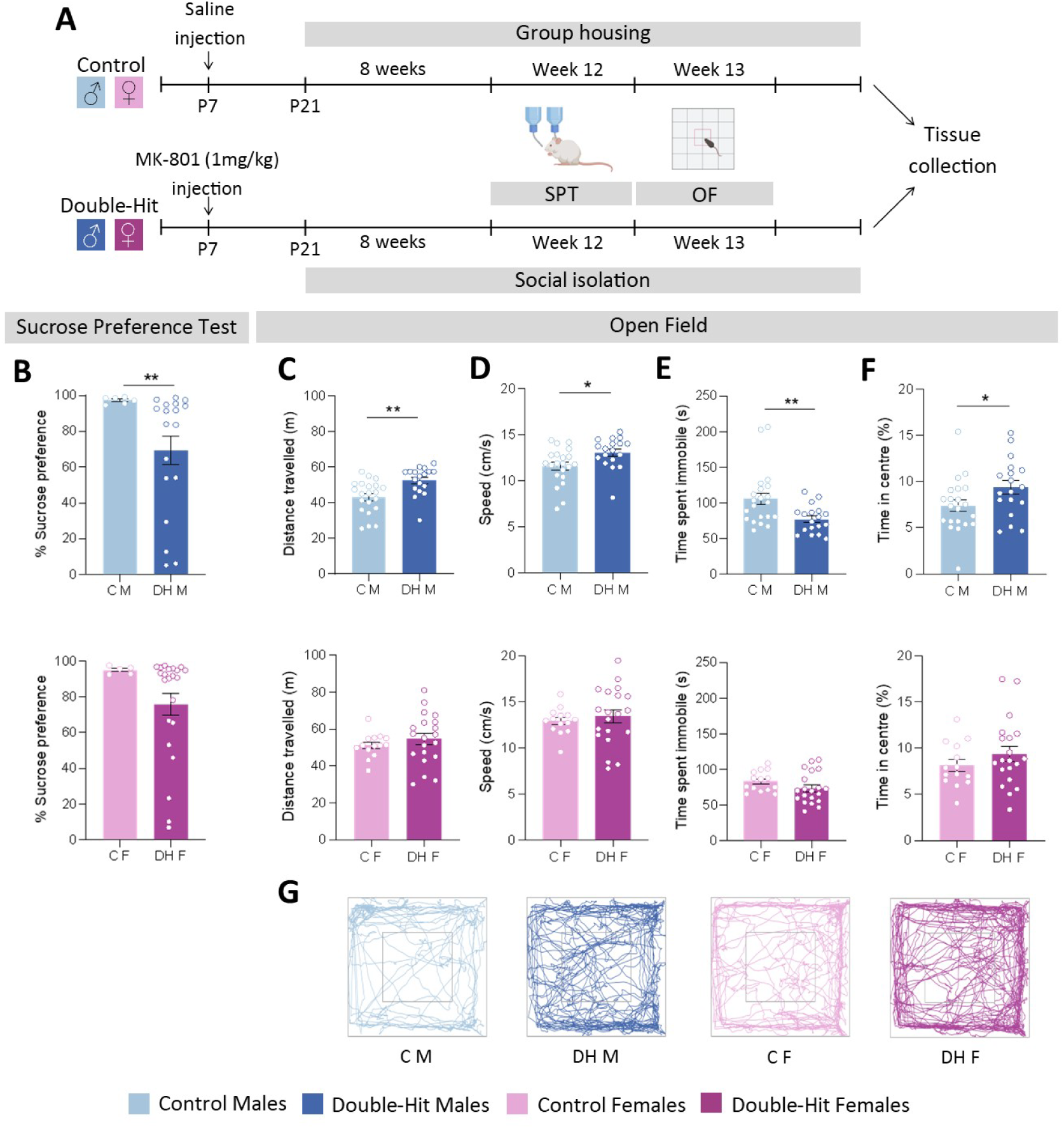
**A** Timeline of the experimental design. Male and female rat pups received intraperitoneally injections of saline or MK-801 at postnatal day 7 (P7). After weaning, control rats were group-housed and double-hit rats were housed individually. Behavioural tests were conducted between weeks 12 and 13, after which brain tissue was colleted for further analysis. **B** Sucrose preference test results. **C-F** parametres measured in open field test: **B** Distance travelled, **C** speed, **D** time spent immobile and E time spent in centre of the arena. **G** Representative movement traces of rats in the open field arena. In all graphs, each dot represents one animal and data is presented as mean ± SEM. **Abreviations**: C M: Control males, C F: control females, DH M: double-hit males, DH F: double-hit females.

### 2. Behavioral tests

Behavioral tests were conducted using 21 double-hit male rats, 24 control male rats, 23 double-hit female rats, and 18 control female rats. Behavioral tests were performed after P90 and included the sucrose preference test, the open field test, and the novel object recognition test. The open field and novel object recognition tests were performed between 8 am and 3 pm in a behavior room separate from the housing facilities.

#### a. Sucrose preference test

On day 1, rats were introduced to two water bottles. On day 2 and 3, they were habituated to a 1% sucrose solution. During habituation, each cage was provided with one bottle containing 1% sucrose solution and one bottle containing water. On day 3 (habituation day 2), the positions of the bottles were switched to minimize side preference. The test was conducted on day 4, during which rats continued to have access to the two bottles, one with water and the other with 1% sucrose solution, for 24 hours. To control for side bias, the position of the bottles was randomly assigned for each cage. Fluid consumption was measured after 24 hours, and sucrose preference was calculated as the volume of 1% sucrose solution consumed divided by total fluid consumption.

#### b. Open field

The testing apparatus was 70x70cm wide and 40 cm tall. Rats were placed in the center of the open field and allowed to explore for 10 minutes. Between each rat, the arena was cleaned with 70% ethanol. Behavior was recorded using an overhead Sony color video camera SSC-E Series and later analyzed using DeepLabCut (Mathis et al., 2018) and a custom script in R (R Core Team, 2024). The behaviors analyzed included total distance traveled, average speed, percentage of the time spent in the center of the open field, and time spent immobile.

### 3. RNA-seq analysis

20 rats were used for the RNAseq analysis (n = 5 per group). Rats were intraperitoneally injected with a high dose of pentobarbital sodium (Dolethal, Vetoquinol) and sacrificed by decapitation using a guillotine. The brains were extracted from the cranium, and the hemispheres were separated. The left hemisphere was frozen in liquid nitrogen and stored at -80°C. The medial prefrontal cortex (mPFC) of the right hemisphere, encompassing both the prelimbic and infralimbic areas, was microdissected and preserved in RNA Stabilization Solution (Invitrogen). Total mRNA was extracted using NucleoZol (Takara Bio Company) according to the manufacturer’s instructions. The concentration and purity of the mRNA were determined using a NanoDrop spectrophotometer and sequenced at the Centro Nacional de Análisis Genómico (CNAG) in Barcelona, Spain.

Read quality was assessed using FastQC (v. 0.11.9). Reads were aligned to the rat reference genome (Rattus_norvegicus.mRatBN7.2) using HISAT2, and read counts were quantified using HTSeq (v. 2.2.1) using a strict intersection with the corresponding Ensembl gene annotation file. (Rattus_norvegicus.mRatBN7.2). Differential gene expression (DEG) analysis was performed using DESeq2 (v. 1.36) implemented in R. Data from female and male rats were analyzed both jointly, including sex as a covariate, and separately. The Benjamin-Hochberg method was used to correct for multiple testing, and genes with an adjusted p-value < 0.05 were considered differentially expressed.

Gene Set Enrichment Analysis (GSEA) was performed using the gsea function in clusterProfiler (v. 4.4.4) in R. Enrichment analyses were conducted using the Gene Ontology (GO), Kyoto Encyclopedia of Genes and Genomes (KEGG), and Human Phenotype Ontology (HPO) databases. The Benjamin-Hochberg method was used to correct multiple testing and obtain the enriched categories defined as those with an adjusted p-value < 0.05.

Multi-marker Analysis of GenoMic Annotation (MAGMA) was performed using data from the schizophrenia GWAS meta-analysis conducted by the Psychiatric Genomic Consortium (67,390 cases and 94,015 controls). Data visualization was obtained using the GOBubble function in GOPlot (v. 1.0.2) in R (Figure 2A).

**Figure 2:**
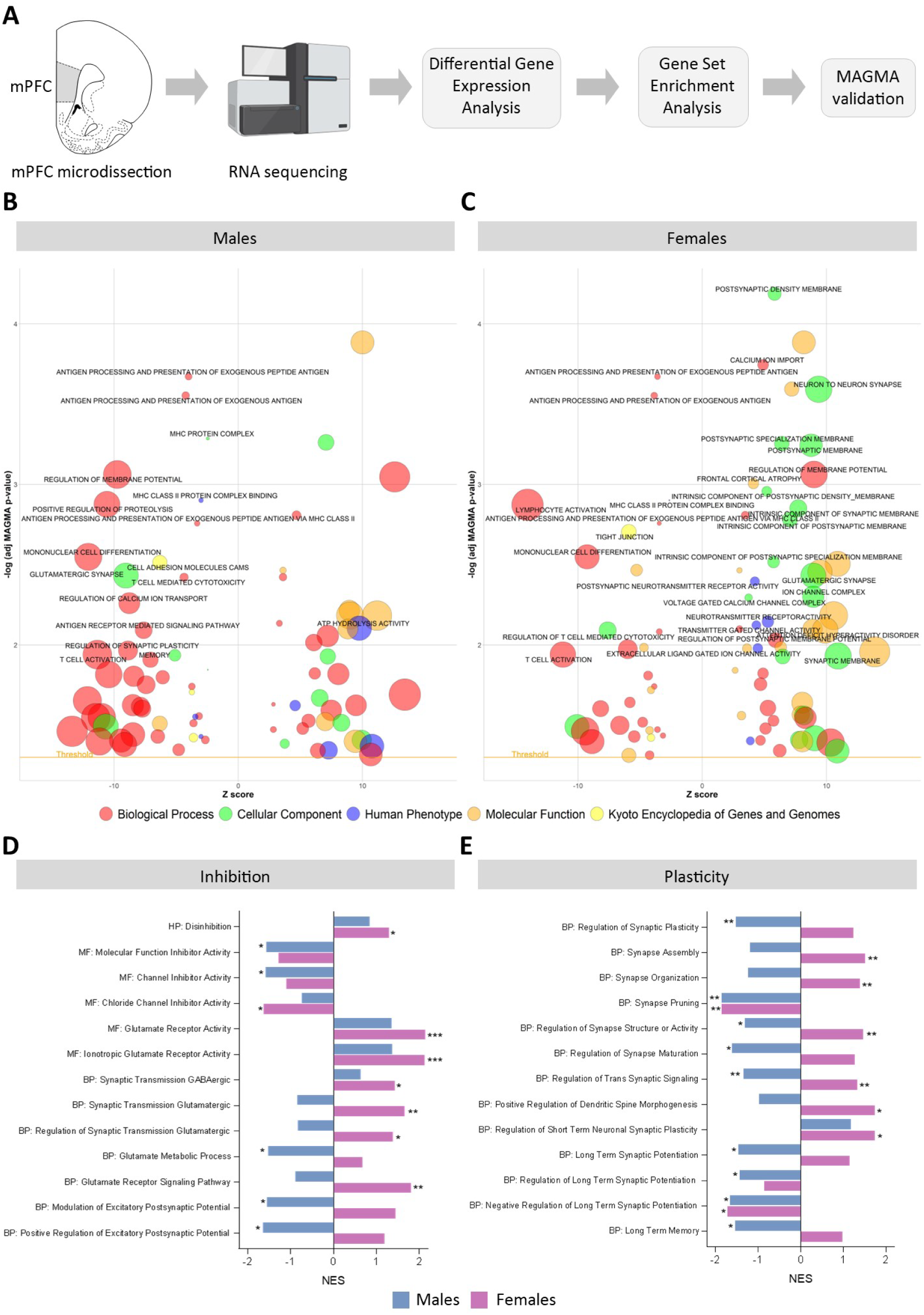
**A** Schematic representation of the steps in the RNAseq analysis pipeline. **B-C** Bubble plots showing the most relevant MAGMA-validated GSEA categories in males **(B)** and females **(C)**. **D-E** GSEA categories specifically associated with inhibition (**D**) and plasticity (**E**) in male and female double-hit rats. **Abreviations**: HP: Human Phenotype, MF: Molecular Function, BP: Biological Process.

To focus on biologically relevant pathways related to inhibition and plasticity, we filtered the GSEA results by selecting categories that contained the following keywords: “synaptic,” “GABA,” “plasticity,” “inhibitory,” “long,” “dendritic,” “neuronal,” “neurotransmitter,” “NMDA,” “AMPA,” “inhibit,” “parvalbumin,” “interneuron,” “neurogenesis,” “neural,” “neuronal,” “synapse,” “action potential,” or “glutamate“. For this targeted, hypothesis-driven analysis, we used nominal *p*-values to identify meaningful patterns within a limited subset of gene sets. Categories were retained for further interpretation only if they showed nominal significance (*p* < 0.05) in either males or females and were functionally relevant to the biological processes of interest.

### 4. Immunofluorescence

41 rats were perfused transcardially with 4% paraformaldehyde in phosphate buffer (PB, 0.1 M, pH 7.4). Brains were extracted from the cranium and post-fixed in the same 4% paraformaldehyde solution for 24 hours. Afterwards, the hemispheres were separated, cryoprotected in a 30% sucrose solution, and stored at -20°C in an antifreeze solution. The left hemispheres were later sectioned at 50 μm in coronal sections using a sliding microtome. These sections were used for immunofluorescence analysis.

Immunofluorescence was performed on free-floating sections as follows: Sections were washed in PBS with 1% Triton X-100 and then incubated for 15 minutes in an antigen retrieval solution (0.01M citrate buffer, pH 6) at 80°C. After additional washes in PBS with 1% Triton X-100, sections were incubated for 1 hour at room temperature in PBS with 1% Triton X-100 and 10% normal goat serum (NGS) to block nonspecific binding. Sections were then incubated overnight at 4°C with the primary antibodies: PV (guinea pig, 1:1.000, Synaptic Systems), Otx2 1:100 (mouse, 1:100, College de France, in-house), biotinylated Wisteria floribunda agglutinin (1:200, Vectorlabs), GAD67 (mouse, 1:1.600, Sigma-Aldrich), γH2A.X (rabbit, 1:2.500, Abcam), CaMKII (mouse, 1:500, Abcam), and cFos (rabbit, 1:2.000, Abcam) diluted in PBS with 1% Triton-X-100 and 3% NGS. The next day, after washing, sections were incubated with the corresponding secondary Alexa Fluor-conjugated antibodies (1:200, Thermofisher) in the same incubation solution. Finally, sections were mounted using Fluoromount-G mounting medium, either with DAPI or without DAPI (Invitrogen), depending on the experimental condition.

#### a. Quantification of immunofluorescence markers

Images were acquired using a Leica Stellaris SP8 confocal microscope with either a 20x or a 40x objective. Separate images were captured for the prelimbic and infralimbic regions of each sample. Fluorescence intensity of each marker was quantified in individual cells using custom macros using FIJI/ImageJ software. Fluorescence of Otx2, GAD67, and γH2AX was measured exclusively in PV+ cells, while cFos fluorescence was measured in CaMKIIα+ cells.

### 5. Electrophysiology

Electrophysiological recordings were performed on 19 adult Lister Hooded rats of both sexes weighing 430-500 g (males) and 260-310 g (females). Animals were anesthetized with urethane (1.6 g/kg intraperitoneally) and placed in a David Kopf stereotaxic apparatus (Tujunga, CA, USA). Supplemental doses of urethane (0.5 g/kg i.p.) were given to maintain areflexia. A midline scalp incision was made, and the periosteum was removed to expose the skull. Small craniotomies were drilled over the medial prefrontal cortex (mPFC) and the basolateral amygdala (BLA).

Tungsten microelectrodes (5 MΩ) were used to obtain single-unit recordings in the prelimbic (PL) or infralimbic (IL) cortical areas of the mPFC (A: +2.8 mm, L: 0.5 mm from the bregma; H: 3.5/4.0 or 4.5 mm, respectively, from the cortical surface), according to the atlas of Paxinos and Watson (2007) (Figure 6A). The unit recordings were filtered between 0.3 and 3 kHz and amplified using a DAM50 preamplifier (World Precision Instruments, Friedberg, Germany). The signals were sampled at 10 kHz through an analog-to-digital converter (Power 1401 data acquisition unit, Cambridge Electronic Design, Cambridge, UK) and fed into a PC for offline analysis with Spike 2 software (Cambridge Electronic Design).

Electrical stimulation was performed by a bipolar stimulation electrode (World Precision Instruments, Friedberg, Germany) aimed at BLA (A: -2.3 mm, L: 8 mm from the bregma; H: 9 mm) by pulses of 0.3 ms duration and <150 μA of intensity delivered at 0.5 Hz. For comparison, the stimulation intensity was set two times higher than the threshold to elicit spike firing in the mPFC neurons. Pair-pulses at 50 or 300 ms delay were applied to test feedback inhibition.

#### a. Drugs

The selective GABAA receptor antagonist bicuculline (10 mM in 0.9% NaCl, pH 3, Sigma, St. Louis, MO, USA) or the GABAB receptor antagonist saclofen (10 mM in 0.9% NaCl, pH 3, Sigma, St. Louis, MO, USA) was applied. Drugs were slowly delivered through a cannula connected to a Hamilton syringe (1 μl) over a one-minute period; recordings were performed 10 min after drug application.

The peristimulus time histograms (PSTHs) were used to calculate spike responses in a 50 ms post-stimulus time window following each electrical stimulus (1 ms bin-width). Pair-pulse stimulation was used to quantify feedback inhibition; the reduction of the response in the second stimuli was calculated.

### 6. Statistics

Statistical analysis of behavioral tests, electrophysiology, and immunofluorescence analysis was performed using GraphPad Prism 9 software. Data from males and females were analyzed separately, except for electrophysiology, where results from both sexes were pooled. Normality and homoscedasticity were assessed, and either *t*-tests or Mann–Whitney tests were applied as appropriate. For graphical representation, mean ± SEM was used in all cases.

Pairwise correlations between behavioral and immunofluorescence variables were assessed using Pearson’s correlation coefficient in R (R Core Team, 2024) using the Hmisc package. Heatmaps of correlation coefficients were generated using reshape2 for data transformation and ggplot2 for visualization.

## Results

### Male double-hit rats showed traits similar to the symptoms suffered by schizophrenic patients

Among the characteristics frequently seen in animal models of schizophrenia that mimic the symptoms of the illness are hyperlocomotion, anxiety, and anhedonia (Enomoto 2007; Barnes, 2014). In order to identify sex differences, we assessed a number of those traits in our model.

The male double-hit model of schizophrenia rats exhibited anhedonia-like behavior in the sucrose preference test when they were tested about the consumption of sweet water. The 1% sucrose solution was less preferred by double-hit rats than by control rats when examining males (**p = 0.0044). Females showed a similar pattern, although it was not statistically significant (p = 0.099) (Figure 1B). Additional details are shown in Supplementary Figure S1C.

Male double-hit rats showed increased distance travelled in the arena (**p = 0.0019), speed (*p = 0.017), and decreased time immobile (**p = 0.0074), which indicates hyperlocomotion/hyperactivity. The same trend was found in females for all measurements: distance travelled (p = 0.39), speed (p = 0.58), and time immobile (p = 0.16), but without statistical significance (Figure 1C-E). Male double-hit rats also showed an increased percentage of time spent in the center of the open field (*p = 0.049). Since hyperlocomotion also suggests a greater utilization of the central area of the arena, we could not conclude that it is a hypoanxiety-like behavior. The same tendency was seen in females, but again without statistical significance (p = 0.27) (Figure 1F).

In summary, the male double-hit rat model of schizophrenia displayed clear behavioral alterations consistent with core symptoms of the disorder, including anhedonia, evidenced by a significantly reduced preference for sucrose solution, and hyperlocomotion, shown by increased distance traveled, higher speed, and reduced immobility in the open field test. Males also spent more time in the center of the arena, although this was likely related to their increased activity rather than reduced anxiety. Female rats exhibited similar behavioral trends across all measures, but these differences did not reach statistical significance, suggesting that the behavioral effects of the double-hit model are more pronounced in males.

## RNA-seq reveals sex-specific dysregulation of synaptic plasticity and E/I balance

To explore the molecular mechanisms underlying the double-hit model of schizophrenia and to assess potential sex differences, RNA-seq was performed on mPFC samples from male and female rats of control and double-hit groups (5 each). This analysis enabled the identification of differentially expressed genes and altered biological processes associated with the pathophysiology of the disorder (Figure 2A).

When analyzing females and males together, a total of 19,538 genes were expressed in the mPFC of the subjects. Of these genes, 3 genes were differentially expressed (DEGs) (adjusted p-value < 0.05) between control and double-hit rats (Supplementary Figure S2A). Two lncRNAs (ENSRNOG00000069843, ENSRNOG00000066269) and *Ch25h*, which codes for the *Cholesterol 25-hydroxylase*, which is involved in the regulation of the interleukin-1β (Supplementary figure S2A).

When analyzing females separately, a total of 19,781 genes were identified, of which 286 were DEGs (adjusted p-value < 0.05) between the control female and the double-hit female rats (Supplementary Figure S2C). Among these, 87 DEGs were seen to be associated with schizophrenia by MAGMA, including glutamate receptors *Grm3* (log2_FC = -0.31, p adj. = 0.043) and *Grinb2* (log2_FC = 0.29, p adj. = 0.043), DNA methyltransferase *Dnmt3a* (log2_FC = 0.25, p adj. = 0.039), histone deacetylase *Hdac4* (log2_FC = 0.31, p adj. = 0.029), neurotrophin receptor *Ntrk3* (log2_FC = 0.30, p adj. = 0.026), and *Cpeb1* (log2_FC = 0.37, p adj. = 0.0014) (Supplementary Figure S2D). The top 10 DEGs could strongly differentiate between control and double-hit female rats (Supplementary Figure S2E). In males, a total of 18,751 were identified, but no DEGs were identified after multiple test corrections (Supplementary Figure S2B).

Gene Set Enrichment Analysis (GSEA) was conducted for the HPO, KEGG, GOCC, GOMF, and GOBP databases for the genes identified when females and males were analyzed together. 1,193 enriched categories were found, of which 110 had significant association with schizophrenia by MAGMA (Supplementary Figure S2D).

In males, of the 783 enriched categories, 80 were found to have significant associations with schizophrenia by MAGMA. Including categories related to antigen processing and presentation, regulation of membrane potential, and regulation of synaptic plasticity (Figure 2B). In females, of the 674 enriched categories, 83 were found to have significant associations with schizophrenia by MAGMA. Including categories related to antigen processing and presentation, postsynaptic membrane, and glutamatergic synapse (Figure 2C).

An analysis of the GSEA was conducted in the categories associated with “brain plasticity” and “inhibition” that were found enriched either for males or for females. For the categories related to “inhibition,” the GSEA identified a shared molecular signature for male and female rats, marked by an upregulation of glutamate receptor activity (MF: glutamate receptor activity (females ***p = 0.000084, males p = 0.12) and MF: ionotropic glutamate receptor activity (females ***p adj = 0.000075, males p = 0.11)) alongside a downregulation of inhibitory signaling pathways (MF: molecular function inhibitory activity (females *p = 0.019, males 0.14), MF: chloride channel inhibitor activity (females *p = 0.020, males p = 0.82), and MF: channel inhibitor activity (males *p = 0.031, females p = 0.29)). We also found an upregulation in genes involved in the HPO category disinhibition, which was significant in females (*p = 0.015). These findings are markers of an E/I imbalance.

However, we can also see sex-specific differences in some biological process categories. In males, we observed a significant downregulation of gene sets involved in glutamate metabolism (BP: glutamate metabolic process (*p = 0.026)) and excitatory postsynaptic modulation (BP: modulation of excitatory postsynaptic potential (*p = 0.022), BP: positive regulation of excitatory postsynaptic potential (*p = 0.015)), suggesting a potential dampening of excitatory neurotransmission despite an upregulated receptor activity. In females, we found a significant upregulation of glutamatergic (BP: synaptic transmission glutamatergic (*p = 0.0022), BP: regulation of synaptic transmission glutamatergic (*p = 0.034), BP: glutamate receptor signaling (*p = 0.0012)) and GABAergic synaptic transmission (BP: synaptic transmission GABAergic (*p = 0.042)). This points to a state of enhanced and possibly dysregulated synaptic activity. Notably, in all these categories, the opposite non-significant trend was observed in the other sex, indicating possible sex-specific susceptibility or compensatory mechanisms (Figure 2D).

For the GSEA categories related to “plasticity,” we found fewer shared mechanisms between males and females. Both sexes showed a significant downregulation of synapse pruning (males **p = 0.0054, females **p = 0.0036) and negative regulation of long-term synaptic potentiation (males *p = 0.027, females *p = 0.012). In males, we found a downregulation of synaptic plasticity mechanisms (BP: regulation of synaptic plasticity (**p = 0.0024), BP: regulation of trans synaptic signaling (**p = 0.0045), BP: regulation of synapse structure or activity (*p = 0.023), and BP: regulation of synapse maturation (*p = 0.028)). Double-hit male rats also showed a downregulation of BP: long-term memory (*p = 0.021). On the contrary, females showed an upregulation of categories governing synaptic plasticity (BP: synapse organization (**p = 0.0019), BP: synapse assembly (**p = 0.0016), BP: regulation of trans-synaptic activity (**p = 0.0060), BP: regulation of synapse structure or activity (**p = 0.0014), BP: regulation of short-term neuronal synaptic plasticity (*p = 0.018), and BP: positive regulation of dendritic spine morphogenesis (*p = 0.014)). These results suggest that both male and female double-hit rats experience a dysregulation of synaptic plasticity, but the direction of these molecular changes is sex dependent. Male double-hit rats seem to exhibit more suppressed or inefficient synaptic plasticity mechanisms, while females appear to have enhanced or dysregulated synaptic plasticity mechanisms (Figure 2E).

Overall, these results reveal a clear sex-specific transcriptional response to the double-hit model of schizophrenia. Female rats exhibited a greater number of altered genes and pathways, particularly those related to glutamatergic and GABAergic neurotransmission, while males showed a suppression of synaptic plasticity mechanisms and glutamate metabolism. These findings indicate a shared E/I imbalance across sexes but with opposing molecular directions.

### The mPFC of the male and female double-hit model of schizophrenia exhibits opposite indicators of inhibition and plasticity

To further characterize the cellular mechanisms underlying the behavioral and molecular alterations observed, we examined markers related to inhibitory plasticity and excitatory-inhibitory balance in the mPFC (Figure 3A, Figure 4A-C).

**Figure 3:**
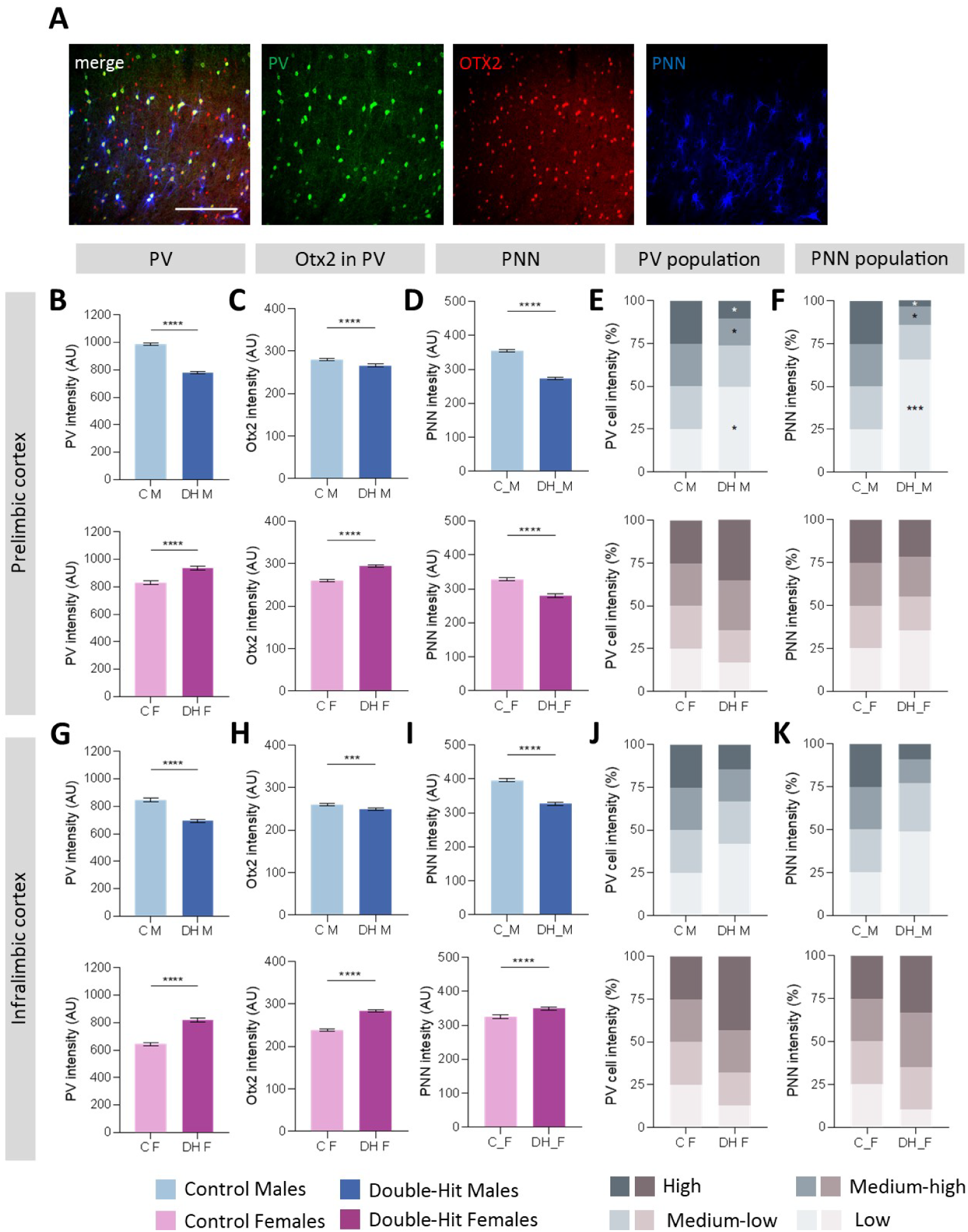
**A** Representative confocal microscopy images from prelimbic (PL) cortex showing parvalbumin (PV) (green), Otx2 (red), and perineuronal net (PNN) (blue) labelling. Images were acquired using a 2Ox objective. **B-D** Intensity of PV (B), Otx2 (**C**) and PNN (**D**) labelling in the prelimbic cortex. **E-F** Distribution of PV+ interneurons (**E**) and PNNs (**F**) according to labelling intensity in the PL cortex. G-l Intensity of PV (**G**), Otx2 (**H**) and PNN (**í**) labelling in the infralimbic (IL) cortex. **J-K** Distribution of PV+ interneurons (J) and PNNs (**K**) according to labelling intensity in the IL cortex. In all graphs, data is presented as mean ± SEM. **Abreviations**: C M: Control males, C F: control females, DH M: double-hit males, DH F: double-hit females, PNN: perineuronal nets. Scale bar: 200 µm.

**Figure 4:**
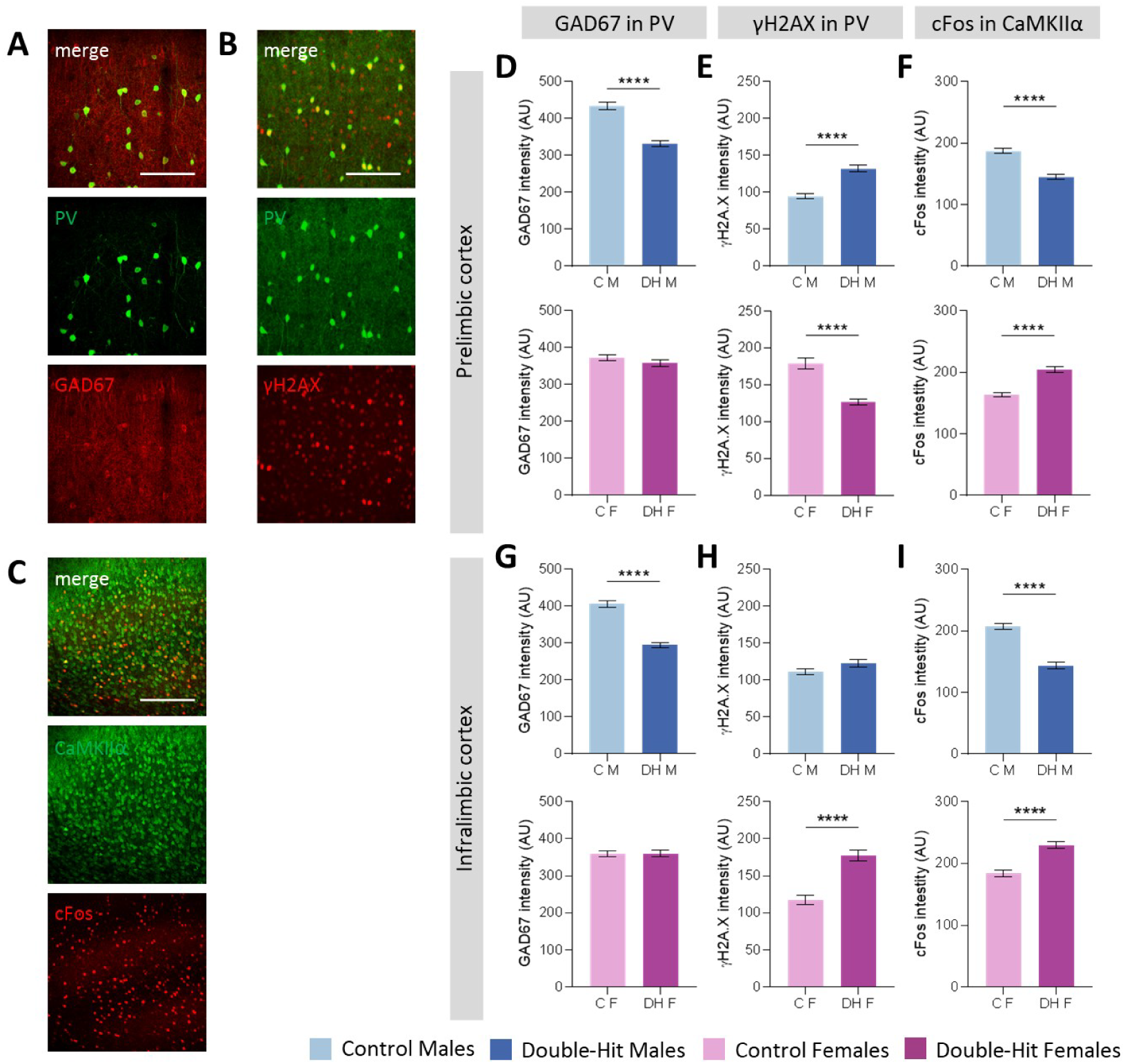
**A-C** Representative confocal microscopy images from prelimbic (PL) cortex showing: **A** PV (green) and GAD67 (red) labeling; **B** PV (green) and γH2AX (red) labeling; C CaMKIIα (green) and cFos (red) labeling. Images were captured using either a 4Ox (**A, B**) or a 2Ox (C) objective. **D-F** Quantification of la veiling intensity in the prelimbic (PL)cortex: GAD67 (**D**) and yH2AX (**E**) in PV+ interneurons and cFos labelling in CaMKIIα+ cells (**F**). **G-l** Quantification of labeling intensity in the infralimbic (IL) cortex: GAD67 (**G**) and yH2AX (**H**) in PV+ interneurons and cFos labelling in CaMKIIα+ cells (I). In all graphs, data is presented as mean ±SEM. Abreviations: C M: Control males, C F: control females, DH M: double-hit males, DH F: double-hit females. Scale bar: 100 µm (A, B); 200 µm (C)

Analyses of PV, OTX2, and PNNs markers were first conducted to determine PV+ interneuron plasticity in the PrL and IL (Figure 3A). We found a decrease of PV labeling intensity in male double-hit rats both in the PrL (****p < 0.0001) (Figure 3B) and IL cortex (****p < 0.0001) (Figure 3G). This change is caused by an increase in low-intensity PV cells and a decrease in high-intensity and medium-high PV cells (Figure 3E, J). These differences were significant in the PrL cortex , but not in the IL cortex. On the contrary, an increase in PV labelling intensity was found in female double-hit rats in both the PrL (****p < 0.0001) (Figure 3B) and IL cortices (****p < 0.0001) (Figure 3G). Nevertheless, there were no significant differences in the distribution of PV cell populations (Figure 3E, J).

In tandem with the PV alterations, we observed that OTX2 intensity, a marker of cell maturation, was decreased in male double-hit rats in the PrL (****p < 0.0001) (Figure 3C) and IL (****p = 0.0001) (Figure 3H), while in female double-hit rats, it increased in the PrL (****p < 0.0001) (Figure 3C) and IL (****p < 0.0001) (Figure 3H).

Additionally, we reported that male double-hit rats had lower PNN intensity in the PrL (****p < 0.0001) (Figure 3E) and IL (****p = 0.0001) (Figure 3I), while female double-hit rats had lower PNN intensity only in the PrL (****p = 0.0001) (Figure 3E) and higher in the IL (****p = 0.0001) (Figure 3I). In the case of the PrL cortex of male double-hit rats, this is brought on by a rise in low-intensity PV and a fall in high-intensity PV. The distribution of PV cell populations in the PrL of male and female double-hit rats did not change significantly (Figure 3F, K).

Next, in order to evaluate inhibition in the mPFC, we examined various markers. When we first examined GAD67 in PV+ cells, we discovered that male double-hit rats had lower levels of GAD67 in both the PrL (****p = 0.0001) (Figure 4D) and IL (****p = 0.0001) (Figure 4G). Neither the PrL (p = 0.34) (Figure 4D) nor the IL (p = 0.76) (Figure 3G) showed any differences in female double-hit rats. Additionally, we examined the expression of the marker of DNA damage γH2AX in PV+ cells and discovered that male double-hit rats had higher γH2AX intensity in their PrL (****p = 0.0001) (Figure 4E), but no differences in their IL (p = 0.079) (Figure 4H). On the contrary, we observed a reduction in γH2AX intensity in the PrL (****p = 0.0001) (Figure 4E) and an increase in γH2AX intensity in the IL (****p = 0.0001) (Figure 4H) in female double-hit rats.

To assess excitatory neuron activation, we examined cFos expression in CaMKIIα+ cells. Male double-hit rats showed decreased expression of cFos in the PrL (****p < 0.0001) (Figure 4F) and the IL (****p < 0.0001) (Figure 4I). Conversely, female double-hit rats showed increased cFos intensity in the PrL cortex (****p < 0.0001) (Figure 4F) and IL (****p < 0.0001) (Figure 4I).

We investigated the relationship between the histology markers examined and the outcomes of the open field test. Speed and distance traveled were positively correlated with PV (p = 0.0067), Otx2 (p = 0.0062), PNN (p = 0.019), and c-Fos in CamKII+ (p = 0.021) cells in the prelimbic cortex, and speed and distance traveled were positively correlated with PV, Otx2, and PNN in the infralimbic cortex (Supplementary Figure S3).

In all, our findings indicate marked sex-dependent differences in cortical inhibitory and excitatory regulation in the double-hit model. Male rats exhibited decreased plasticity and impaired inhibitory interneuron function with reduced excitatory activity, while females showed opposite trends suggestive of enhanced or dysregulated cortical activity. These results suggest that the double-hit model induces an imbalance in excitatory-inhibitory mechanisms that manifests differently between sexes, potentially contributing to the distinct behavioral and molecular phenotypes observed.

### Disinhibited cortical activity is observed in both male and female double-hit animal models of schizophrenia

To assess potential functional alterations in cortical inhibitory control in the double-hit model of schizophrenia, we performed in vivo electrophysiological recordings in the PrL and IL regions of the mPFC during basolateral amygdala (BLA) stimulation (Figure 6A). Single- and paired-pulse protocols were applied to evaluate synaptic responsiveness and feedback inhibition and to explore possible sex-dependent differences in cortical excitability.

Units recorded in spontaneous conditions were silent or showed a low firing rate in both control (1.74 ± 0.41 spikes/s; 1.69 ± 0.36 spikes/s in male and female rats, respectively) and double-hit animals (1.82 ± 0.36 spikes/s; 1.88 ± 0.49 spikes/s in male and female rats, respectively) (Supplementary Figure 4A-C).

According to previous findings (Garcia-Mompo et al., 2020), double-hit rats may show a decrease in the cortical feedback inhibition. To test this hypothesis, paired pulses at 50 or 300 ms delays were applied in the BLA while unit recordings were performed in PrL and IL cortical areas. When combining males and females, the response to the second pulse at 50 ms delay was reduced by -12.8 ± 2.5% (n = 75) in PrL and -13.0 ± 5.1% (n = 23) in IL. However, double-hit rats showed paired-pulse facilitation at the same 50 ms delay both in PrL, +14.4 ± 4.5% (n = 53), and in IL, +18.3 ± 5.5% (n = 25). Differences with control were statistically significant at 50 ms delay both in PrL (****p < 0.0001) and in IL (****p < 0.0001) neurons (Figure 5C).

**Figure 5:**
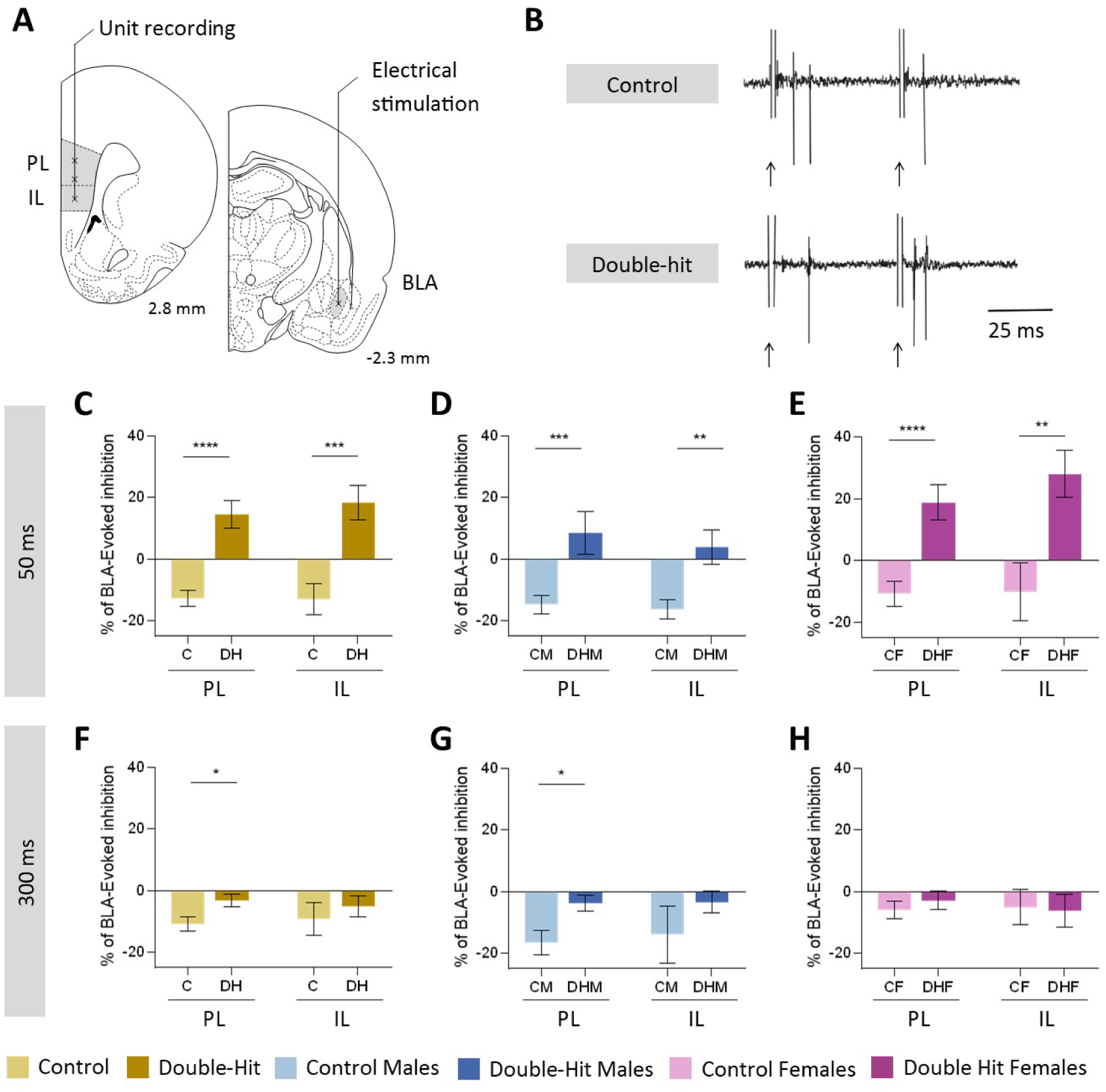
**A** Schematic representation of experimental setup. The stimulation electrode was placed in the basolateral amygdala (BLA) and the recording electrode was placed either in the prelimbic (PL) or infralimbic cortex (IL). **B** Representative response in the PL following stimulation of the BLA (arrows). **C-H** Unit recordings of BLA-evoked inhibition in the PL and IL cortex. Responses are shown when pulses are separated by 50 ms in all rats combined (**C**), males only (**D**) and females only €, and when pulses were separated by 300 ms in all rats combined (**F**), males only (**G**) and females only (**H**). In all graphs, data is presented as mean ± SEM. Abreviations: C M: Control males, C F: control females, DH M: double-hit males, DH F: double-hit females, PL: prelimbic cortex, IL: infralimbic cortex, BLA: basolateral amygdala.

**Figure 6:**
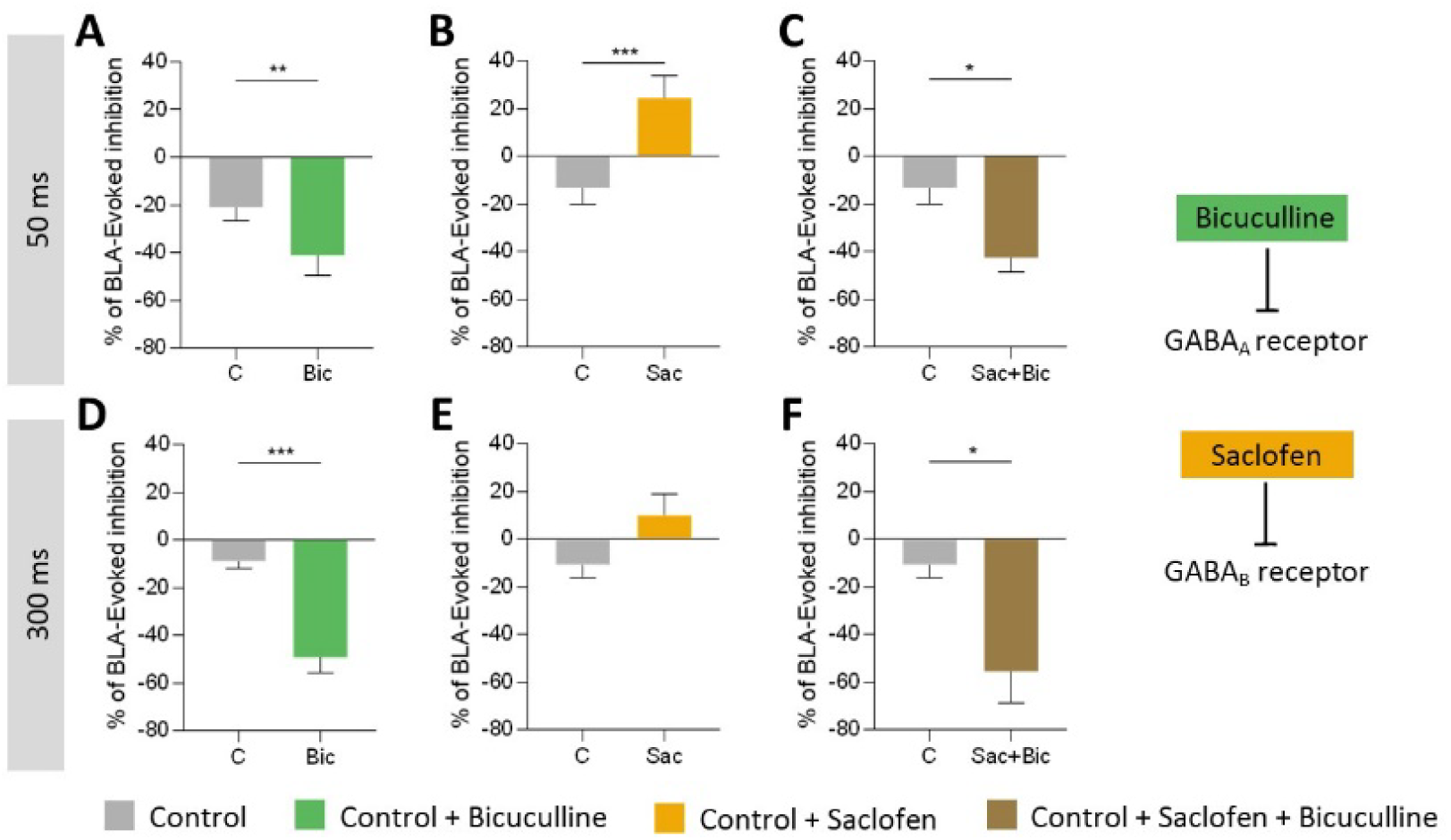
**A-F** Effects of bicucullin, saclofen, or their combination on unit recording of-BLA evoked inhibition in the prelimbic (PL) cortex of male and female control rats combined. Panels show responses after the addition of bicuculline (**A, D**), saclofen (**B, E**), or saclofen followed by biccuculline (**C, F**) with pulses are separated by 50 (**A-C**) or 300 ms (**D-F**). **G** Schematic representatation of the effects of biccuculline and saclofen on GABA receptors. In all graphs, data is presented as mean ± SEM. **Abreviations**: C: control, Bic: bicuculline, Sac: saclofen.

At 300 ms delay, the paired-pulse inhibition was smaller in control rats, -10.8 ± 2.4% (n = 64) in PrL and -9.3 ± 5.3% (n = 23) in IL. Double-hit rats showed decreased pair-pulse inhibition in PrL-3.3 ± 2.0% (n = 55) and in IL -5.1 ± 3.4% (n = 25), which is statistically significant in the case of the PrL cortex (*p = 0.022) but not in the case of the IL (p = 0.51) (Figure 5F).

When sexes were analyzed separately, in control males, the response to the second pulse at 50 ms delay was reduced by -14.7 ± 3.0% (n = 39) in the PrL and by -16.2 ± 3.1% (n = 11) in the IL. However, double-hit male rats showed a paired-pulse facilitation at the same 50 ms delay in both the PrL, 8.6 ± 7.0% (n = 23), and the IL, +3-8 ± 5.5% (n = 10). Differences with male control rats were statistically significant at 50 ms delay both in the PrL (***p = 0.0008) and in the IL neurons (**p = 0.0042) (Figure 5D).

At 300 ms delay, the paired-pulse inhibition was similar in male control rats, -16.6 ± 3.9% (n = 29) in the PrL and -13.9 ± 9.3% (n = 11) in the IL. Male double-hit rats showed reduced paired-pulse inhibition in the PrL, -3.9 ± 2.6% (n = 23), and in the IL, -3.4 ± 3.5% (n = 10). Statistically significant differences were found between male double-hit rats and male rats in the PrL (*p = 0.013), but not in the IL (p = 0.32) (Figure 5G).

In control females, the response to the second pulse at 50 ms delay was reduced by -10.7 ± 4.2% (n = 36) in the PrL and -10.1 ± 9.4% (n = 12) in the IL. However, female double-hit rats showed a paired-pulse facilitation at the same 50 ms delay in both the PrL, +18.8 ± 5.8% (n = 30), and the IL, 28.0 ± 7.6% (n = 15). Differences with control rats were statistically significant at 50 ms delay in the PrL (****p < 0.0001) and in the IL recordings (**p = 0.0037) (Figure 5E).

At 300 ms delay, the paired-pulse inhibition was smaller in female control rats, -6.0 ± 2.9% (n = 35) in the PrL and -5.0 ± 5.7% (n = 12) in the IL. Female double-hit rats showed reduced paired-pulse inhibition in the PrL, -2.9 ± 3.0% (n = 32), and the IL, -6.3 ± 5.3% (n = 15) at 300 ms delay. No statistical differences were found between control and double-hit rats in the PrL (p = 0.45) nor the IL (p = 0.87) (Figure 5H).

The same tendency was found when analyzing field potentials, although statistical significance was only found in the PrL cortex with a 50 ms delay when males and females were analyzed together (***p = 0.0009) and in females (**p = 0.004), and in the IL cortex with a 300 ms delay in males (*p = 0.043) (Supplementary Figure S4D-I).

Overall, the results demonstrate a disruption of cortical inhibitory mechanisms in double-hit rats, characterized by a shift from paired-pulse inhibition to facilitation at short interstimulus intervals. This effect was evident in both PrL and IL and consistent across sexes, although statistical significance was stronger in males. These findings indicate that the double-hit model induces impaired feedback inhibition and enhanced cortical excitability, reflecting a loss of inhibitory tone within prefrontal circuits associated with schizophrenia-like pathology.

### GABAB, but not GABAA, receptors mediate paired-pulse inhibition in the prefrontal cortex

To identify the receptor mechanisms underlying paired-pulse inhibition in the mPFC, we locally applied GABA receptor antagonists in the mPFC of control rats. Specifically, we tested the effects of the GABAA receptor antagonist bicuculline and the GABAB receptor antagonist saclofen, alone and in combination, to determine their respective contributions to cortical inhibitory feedback and to compare them with the electrophysiological alterations observed in the double-hit model.

Contrary to expectations, paired-pulse inhibition was not blocked by local application of the GABA_A_ receptor antagonist bicuculline in control rats (20 mM, 0.2 μl). Bicuculline increased paired-pulse inhibition at 50 ms delay from -20.8 ± 5.9% (n = 9) in basal conditions to -40.9 ± 8.4% (n = 9) after bicuculline application (**p = 0.0057) (Figure 6A). At 300 ms delay, paired-pulse inhibition increased from -8.8 ± 3.2% (n = 11) in basal conditions to 49.0 ± 6.8% (n = 10) after bicuculline application (***p = 0.0003) (Figure 6D).

Thus, we tested the participation of GABA_B_ receptors in the pair-pulse inhibition evoked by BLA stimulation. Local application of the GABA_B_ receptor antagonist saclofen in control rats (20 mM, 0.2 μl) blocked paired-pulse inhibition and caused a facilitation similar to the response obtained in double-hit rats. At 50 ms delay, paired-pulse inhibition changed to facilitation, from -13.3 ±

7.0% (n = 10) in basal conditions to +24.2 ± 9.4% (n = 10) after saclofen application (***p = 0.0005) (Figure 6B). At 300 ms delay, paired-pulse inhibition also changed to facilitation, from 10.8 ± 5.4% (n = 11) in basal conditions to 9.7 ± 9.2% (n = 11 neurons) after saclofen application, but this time without statistical significance (p = 0.12) (Figure 6E).

To study the effect of inhibiting both GABA_A_ and GABA_B_ receptors consecutively, we applied saclofen followed 10 minutes after by bicuculline. After bicuculline application, at 50 ms delay, saclofen induced paired-pulse facilitation changed back to inhibition, from 24.2 ± 9.4% (n = 10) to -42.4 ± 6.4 (n = 6) (*p = 0.034) (Figure 6C). At 300 ms delay, bicuculline application also changed paired-pulse facilitation to inhibition, from 9.7 ± 9.2% (n = 11) to -55.7 ± 13.1% (n = 5) (*p = 0.046) (Figure 6F).

The same tendency was found when analyzing field potentials, although statistical significance was only found in with saclofen at a 50 ms delay (*p = 0.015), and with the combination of saclofen and bicuculline at a 300ms delay (*p = 0.011) (Supplementary figure S5).

These experiments revealed that paired-pulse inhibition in the mPFC is primarily mediated by GABAB receptors rather than GABAA receptors. Contrary to expectations, bicuculline did not block inhibition but instead enhanced it, whereas saclofen abolished inhibition and induced facilitation similar to that observed in double-hit rats. When both antagonists were applied sequentially, bicuculline restored inhibition following saclofen-induced facilitation. Together, these findings indicate that GABAB receptor signaling plays a dominant role in regulating BLA-driven cortical inhibition and that disruption of this mechanism may underlie the impaired inhibitory control seen in the double-hit schizophrenia model.

## Discussion

In this study, we identify sex-dependent molecular and cellular mechanisms that underlie a shared phenotype of mPFC disinhibition in a double-hit model of schizophrenia. Although both males and females ultimately converged on impaired cortical inhibition, the biological routes leading to this dysfunction were strikingly divergent, revealing distinct vulnerabilities in inhibitory circuit maturation and synaptic plasticity.

Male double-hit rats exhibited robust schizophrenia-like behaviors and showed a consistent reduction of PV-, OTX2-, and PNN-related markers, alongside decreased GAD67 and increased γH2AX in PV+ interneurons in accordance with altered inhibition observed previously in this model (Castillo-Gómez et al., 2017; Garcia-Mompo et al., 2020; Gilabert-Juan et al., 2013; Klimczak et al., 2021, 2024). This profile indicates destabilized interneuron maturation, impaired GABAergic transmission, and heightened cellular stress, all of which align with postmortem observations in male patients (Hughes et al., 2024). Transcriptomic analyses further supported this phenotype, showing downregulation of pathways involved in synaptic plasticity, glutamate metabolism, and excitatory modulation. Together, these results point toward a hypoplastic, functionally weakened prefrontal microcircuit.

In contrast, female double-hit rats displayed a markedly different pattern. Despite milder behavioral alterations, females showed increases in PV and OTX2 intensities, higher excitatory neuron activation, and a transcriptomic upregulation of glutamatergic, GABAergic, and synapse assembly pathways. These findings suggest that early-life perturbations in females do not suppress interneuron maturation but instead promote an overactive or dysregulated plasticity state (Santos-Silva et al., 2024; Woodward et al., 2023; Woodward & Coutellier, 2021). Such a mechanism may reflect compensatory responses driven by hormonal or developmental factors and is consistent with human evidence showing that women with schizophrenia often preserve cognitive and social functioning longer than men, yet also present unique neurobiological features (Chi et al., 2024; Moniem & Kafetzopoulos, 2025).

Despite these divergent mechanisms, in vivo electrophysiological recordings revealed a common functional endpoint: a robust shift from paired-pulse inhibition to facilitation following BLA stimulation. This prefrontal disinhibition is also found in both male and female patients of schizophrenia (Radhu et al., 2015; Takahashi et al., 2013). Pharmacological experiments showed that this deficit arises primarily from compromised GABAB receptor-mediated inhibitory feedback, rather than GABAA receptor dysfunction. The ability of saclofen to fully reproduce the double-hit phenotype highlights the centrality of GABAB signaling in maintaining prefrontal inhibitory integrity. These results align with clinical evidence showing altered GABAB receptor expression and function in schizophrenia and position GABAB-dependent circuits as a key node of vulnerability (de Jonge et al., 2017; Fatemi et al., 2011).

The combination of opposite molecular signatures with a shared physiological impairment suggests that cortical disinhibition in schizophrenia may arise through multiple sex-specific mechanisms that ultimately destabilize prefrontal computation. This framework provides a mechanistic explanation for well-established sex differences in disease onset, symptom severity, and progression, while emphasizing the need for sex-informed therapeutic approaches targeting inhibitory plasticity and GABAB-dependent circuit function.

Several limitations should be considered when interpreting these findings. First, although the double-hit model captures key neurodevelopmental features of schizophrenia, it cannot fully recapitulate the complexity and heterogeneity of the human disorder. Second, the reliance on bulk RNA-seq limits the resolution of cell type-specific transcriptional changes, particularly within heterogeneous interneuron populations. Third, electrophysiological experiments were conducted under urethane anesthesia, which may influence baseline cortical excitability and inhibitory tone. Moreover, although sex differences emerged across molecular and histological measures, behavioral alterations in females were mild, potentially reducing the power to detect sex-specific phenotypes at the functional level. Finally, the cross-sectional design precludes determining whether the observed differences reflect developmental delays, compensatory adaptations, or stable pathological states.

In conclusion, our findings reveal that early-life NMDA receptor blockade combined with social isolation produces sex-specific molecular and cellular alterations in the mPFC, yet converges on a common deficit in GABAB receptor-mediated inhibition. Males exhibit impaired interneuron maturation, reduced inhibitory tone, and suppressed plasticity, whereas females show enhanced or dysregulated plasticity and heightened neuronal activation. Both trajectories culminate in cortical disinhibition, a hallmark of schizophrenia.

These results highlight fundamentally different biological pathways leading to similar circuit-level dysfunction and underscore the importance of incorporating sex as a critical variable in mechanistic studies and in the development of targeted interventions for schizophrenia.

Future work should aim to dissect the cell type-specific mechanisms driving sex-dependent trajectories, ideally using single-nucleus or cell-specific transcriptomics to resolve interneuron subpopulations and circuit-level remodeling. Longitudinal studies spanning early development to adulthood will be essential for determining when sex-specific divergence in inhibitory plasticity emerges and whether it can be modulated by hormonal or environmental factors. Combining this approach with optogenetic or chemogenetic manipulation of PV interneurons and GABAB signaling could help establish causal links between molecular alterations and circuit dysfunction. Finally, given the distinct mechanisms observed in males and females, therapeutic strategies should explore sex-informed interventions targeting inhibitory maturation, GABAB-dependent feedback control, and synaptic plasticity pathways, potentially revealing novel avenues for precision medicine in schizophrenia.

## Ethics declarations

The authors declare that they have no conflicts of interest.

The ethics protocol under which the experiments were approved was PROEX 209.7/22. All animal experimentation was conducted in accordance with Directive 2010/63/EU of the European Parliament and of the Council of 22 September 2010 on the protection of animals used for scientific purposes and was approved by the Committee on Bioethics of the Universidad Autónoma de Madrid.

