## Supplemental Figures for "Convergent pathways with impaired inhibition at the frontal cortex are the outcome of differential alterations in the male and female schizophrenia model"

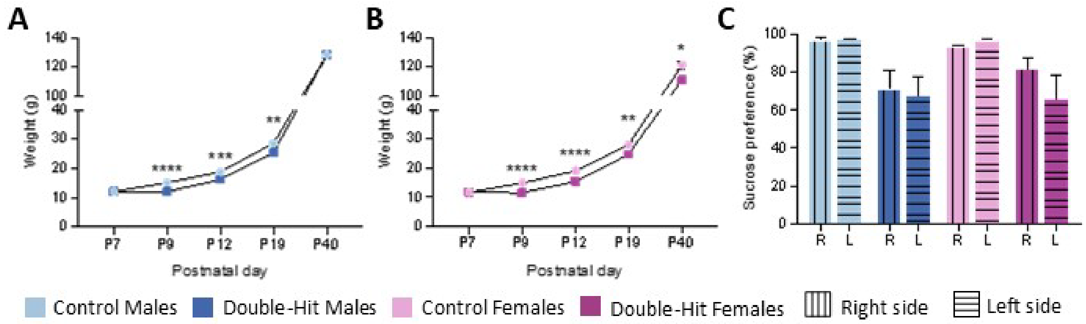


**Supplementary figure 1: A-B** Body weight changes in rats from postnatal day 7 (P7), immediately before MK-801 or saline injection, through postnatal day 40 (P40). **C** Sucrose preference as a function of 1% sucrose bottle position to asses side bias. In all graphs, data is presented as mean ± SEM. **Abreviations:** C M: Control males, C F: control females, DH M: double-hit males, DH F: double-hit females, R: Right side, L: Left side.

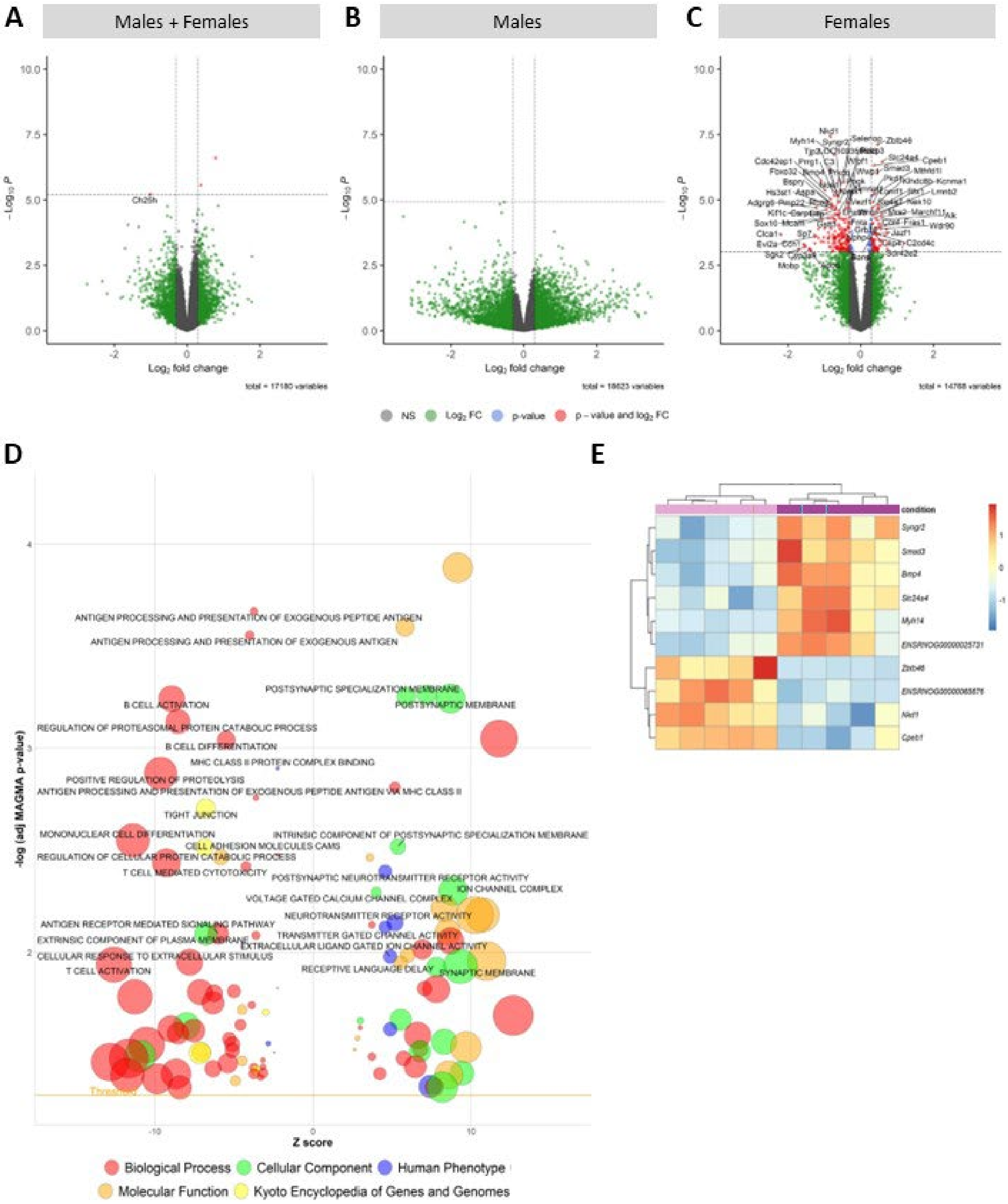


**Supplementary Figure 2: A-C** Volcano plots of differentially expressed genes in males and females together (**A**), males (**B**) and females (**C**). **D** Bubble plot showing the most relevant MAGMA-validated GSEA categories in when males and females are analyzed together. **E** Heatmap of top 10 differentialy expressed genes found in females.

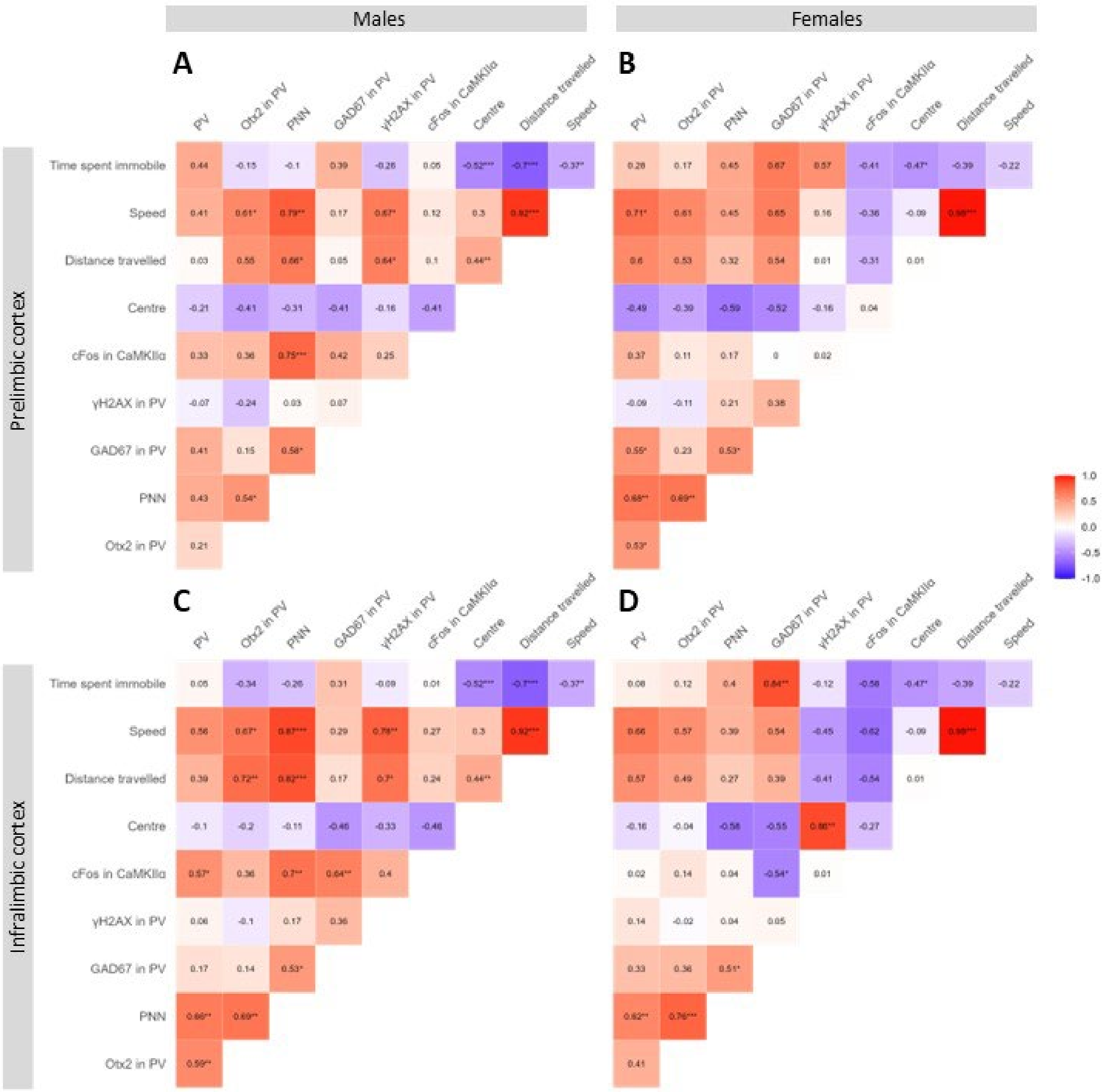


**Supplementary figure 3:** Correlation analysis of inmunostating markers and behavioural parameters in male rats in the PL (**A**) and IL cortex (**B**) and female rats in the PL (**C**) and IL cortex (**D**).

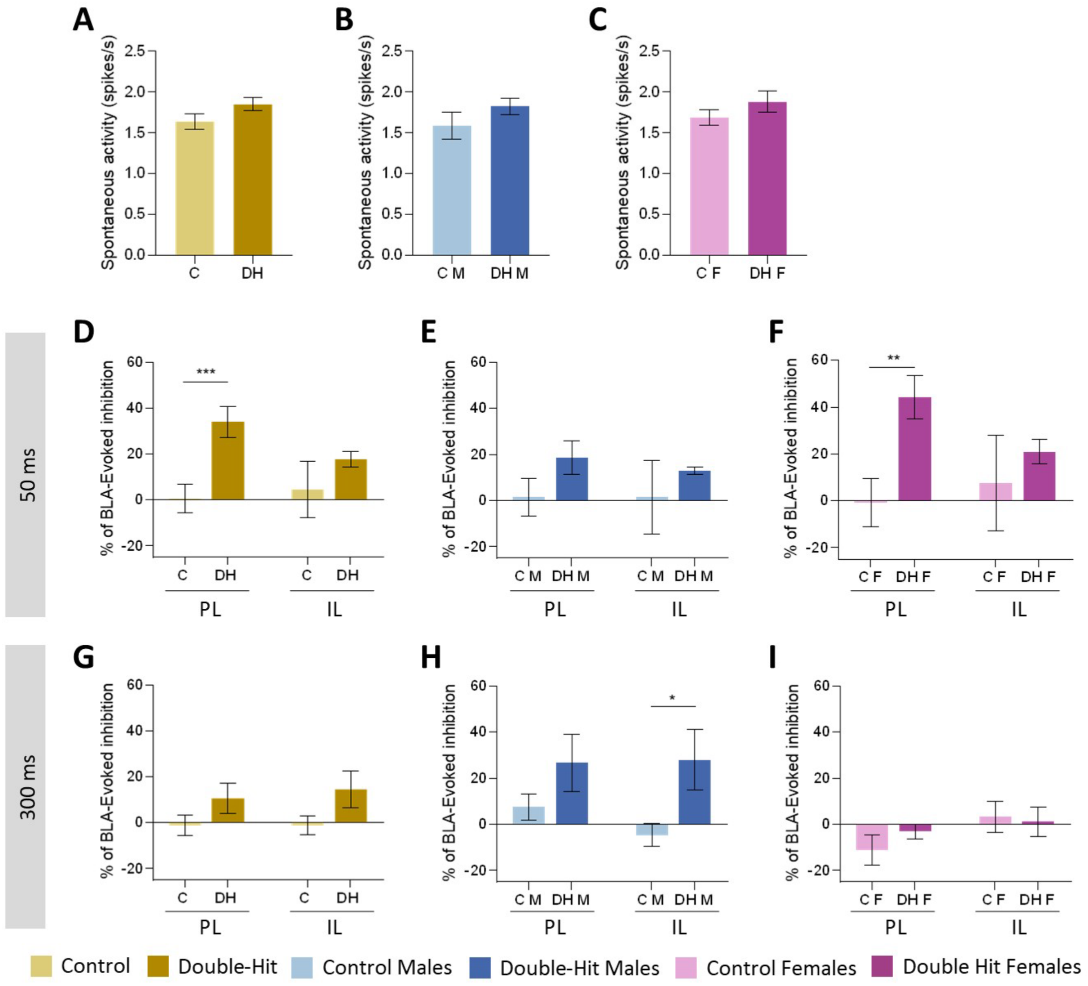


**Suplementary figure 4: A-C** Spontaneous activity in the mPFC in all rats combined (**A**), males only (**B**) and females only (**C**). **D-I** Field activity recordings of BLA-evoked inhibition in the prelimbic (PL) and infralimbic (IL) cortex. Responses are shown when pulses were separated by 50 ms in all rats combined (**D**), males only (**E**) and females only(**F**) and when pulses were separated by 300 ms in all rats combined (**G**), males only (**H**) and females only (**I**). In all graphs, data is presented as mean ± SEM. **Abreviations:** C M: Control males, C F: control females, DH M: double-hit males, DH F: double-hit females, PL: prelimbic cortex, IL: infralimbic cortex, BLA: basolateral amygdala.

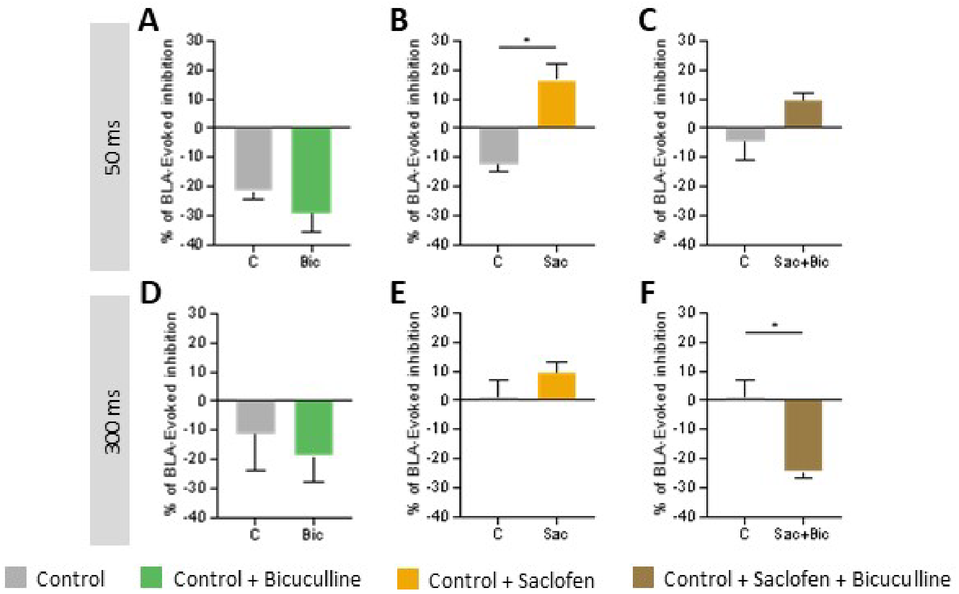


**Suplementary figure 5:** Effect of bicuculline, saclofen, or their combination on field activity recordings of BLA-evoked inhibition in the PL cortex of male and female control rats. Panels show responses after application of bicuculline (**A, D**), saclofen (**B, D**) or saclofen followed by biccuculline (**C, F**) with pulses are separated by 50 (**G-I**) or 300 ms (**J-L**). In all graphs, data is presented as mean ± SEM. **Abreviations:** C: control, Bic: bicuculline, Sac: saclofen.
